# EEG Bad-Channel Detection Using Multi-Feature Thresholding and Co-Occurrence of High-Amplitude Transients

**DOI:** 10.64898/2026.02.04.703874

**Authors:** Amilcar J. Malave, Blair Kaneshiro

## Abstract

Bad channels in electroencephalography (EEG) recordings can substantially degrade downstream analyses, particularly in high-density datasets where localized hardware or motion-related artifacts may affect groups of electrodes in a structured manner. We introduce a MATLAB Module for bad-channel quality control that emphasizes interpretability, relational structure, and human-in-the-loop validation rather than fully automated rejection. The method operates on multichannel EEG data and combines complementary channel-level features, including time-dependent neighbor dissimilarity and amplitude- and variance-based statistics to score and pre-label channels as good, suspicious, or bad. To expose shared artifactual structure, channels are additionally grouped using a similarity measure derived from the co-occurrence of robustly detected high-amplitude transients, allowing channels to be reviewed together. Importantly, clustering is used as an exploratory tool to reveal co-artifactual patterns rather than to impose final class labels, which are confirmed through an interactive review interface supported by summary visualizations and grouped channel displays. This Module is released as a publicly available codebase with documentation, example workflows, and a supporting dataset. This Module is designed as a quality-control step preceding ICA and does not replace end-to-end data cleaning pipelines, which typically involve additional steps such as interpolation of known bad channels.

## Introduction

Electroencephalography (EEG) recordings are sensitive to non-physiological contamination arising from electrode– scalp coupling, motion, and hardware or environmental interference [1]. Even under careful acquisition protocols, it is common for a subset of channels to exhibit persistent failures (e.g., poor contact) or intermittent failures (e.g., transient electrode events or movement-related bursts), particularly in high-density montages and longer sessions [2]. Because these channel-level failures can bias rereferencing and distort multichannel spatial structure, identifying problematic sensors is a standard quality-control step prior to downstream cleaning and analysis steps [1–3].

A variety of automated and semi-automated approaches for EEG bad-channel detection are already widely used, including feature-based thresholding pipelines, correlation- and predictability-based criteria, and artifact-suppression frameworks integrated into common preprocessing toolboxes (e.g., FASTER, PREP, EEGLAB [1–3]). These methods are effective at scale and have become standard components of many workflows. However, they also highlight a persistent practical challenge: channel failure is often time-varying and structured across sensors, and fully automated decisions can obscure why specific channels were flagged, complicating study-specific tolerance decisions and reproducibility. A detailed comparison with existing approaches is provided in the Discussion.

Importantly, not all high-amplitude or transient events observed in EEG recordings indicate defective channels. Transients that recur across multiple electrodes often reflect spatially distributed artifacts or physiological processes (e.g., ocular activity, muscle bursts, motion, or reference-related effects), rather than channel-specific failure. Such shared structure is central to later preprocessing stages, particularly component-based methods such as independent component analysis (ICA), which explicitly rely on cross-channel covariance to identify and remove these artifacts. Prematurely rejecting channels based solely on the presence of shared transients risks discarding physiologically valid information and degrading downstream decomposition performance.

Here we introduce a MATLAB Module for bad-channel quality control that emphasizes interpretable feature summaries and pattern-based grouping, with final decisions confirmed during interactive review. As summarized in Figure 1, the method first computes complementary channel features — including time-dependent neighbor similarity and amplitude/variance-based statistics — to score and prelabel channels as *good, suspicious, or bad*. It then groups channels using an event-timing similarity measure derived from robustly detected high-amplitude EEG transients, enabling channels affected by shared artifact sources to be reviewed together. Importantly, clustering is used to expose co-artifactual structure rather than to impose automatic class labels: final decisions are made through an interactive review step supported by summary visualizations and grouped channel displays. The Module is intended to operate on EEG data that have already undergone filtering and possibly removal of segments not corresponding to analytic epochs. It is designed as a quality-control step preceding ICA and does not replace end-to-end data cleaning pipelines, which typically involve additional steps such as interpolation of known bad channels. This ordering ensures that channel-specific failures are addressed while preserving other shared transient structures, such as eye blinks, that are more appropriately handled by ICA.

**Fig. 1.**
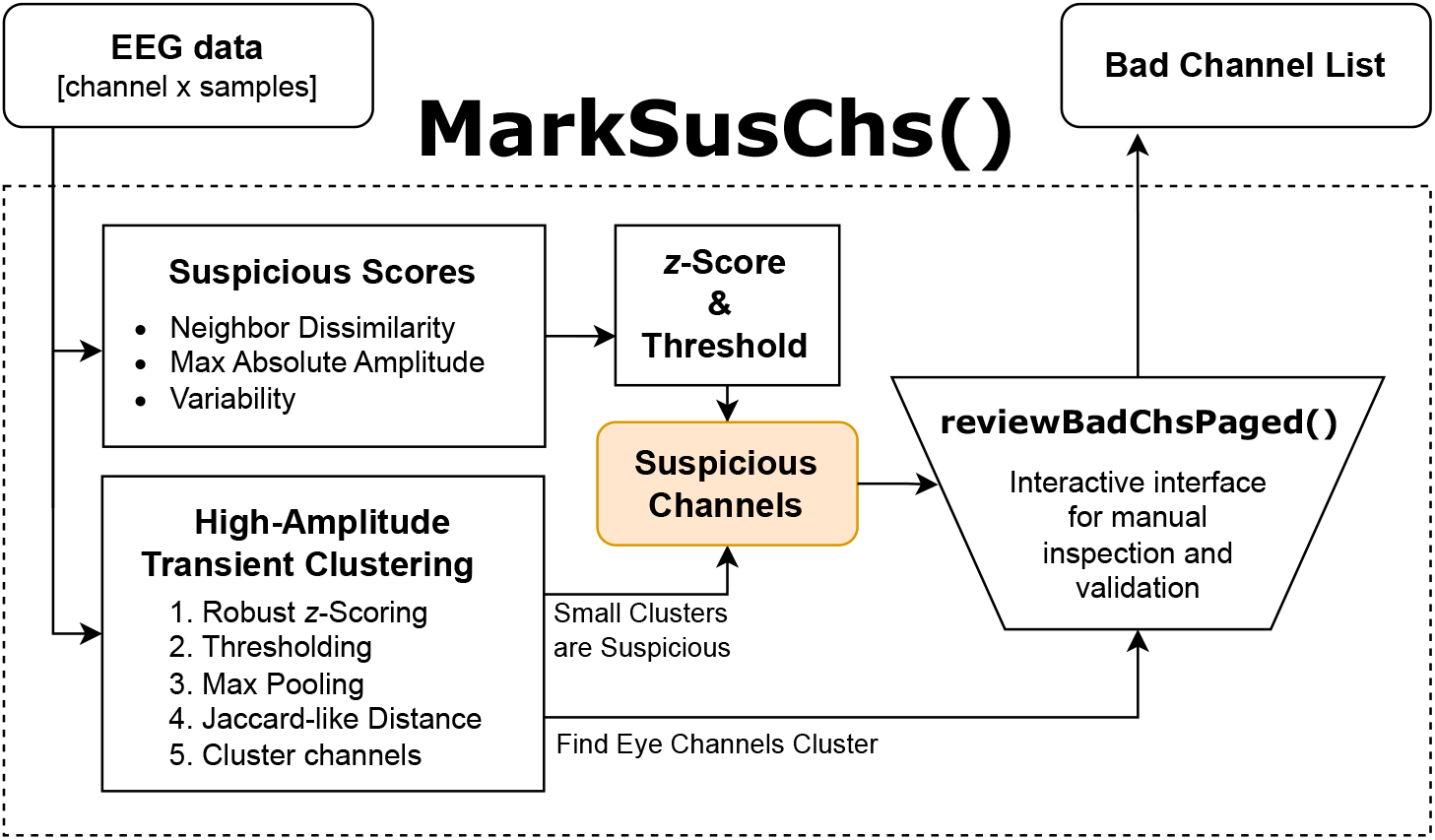
Bad-Channel Detection Module overview. The workflow computes multiple channel-level features, derives an event-timing representation from robust high-amplitude transients, clusters channels by event-timing similarity, and supports a manual review that returns the final bad-channel list.

The project is released as a self-contained and documented Module, with an emphasis on transparency, interpretability, and human-in-the-loop validation. The initial release includes three main components:

1. A publicly available GitHub repository containing the complete codebase, documentation, and usage examples [4].^1^
2. A standalone example dataset used for the illustrative analyses, with accompanying scripts for automated download and local setup [5].^2^
3. This preprint, which describes the methodological rationale and design of the Module.

The remainder of this preprint is organized as follows. We describe the feature definitions, event-based channel similarity metric, and clustering procedure. We then demonstrate the workflow on representative EEG recordings to illustrate how shared artifact structure is revealed and how human validation refines automated suggestions. We close with a discussion of scope, limitations, and intended use within broader EEG preprocessing pipelines.

## Codebase

### Code access and license

The SENSI EEG PREPROC Bad-Channel Detection Module is publicly available as a GitHub repository (see footnote 1). This preprint corresponds to the current tagged GitHub release of the Module. The Module is released under the MIT license.^3^

In addition to the codebase, the release includes a standalone example dataset hosted on the Stanford Digital Repository (SDR) (see footnote 2): the example script in the repository (`example.m`) automatically downloads the SDR files into the local ***ExampleData*** folder.

If using the Module (code, figures, or methods), please cite (i) the GitHub repository, (ii) this preprint, and (iii) the example dataset when used for demonstrations or external projects.

### Dependencies and MATLAB versions

The Module requires a MATLAB^4^ license and depends on one MathWorks toolbox: Statistics and Machine Learning Toolbox.^5^ The Module was developed and tested on MATLAB R2024b. It was validated on Windows and Linux systems; we anticipate that it will work across operating systems, but this has not been comprehensively validated.

### Folders and files in the repository

The top level of the repository contains the following directories and files:

#### EGIMontages

Contains montage-specific neighbor-electrode maps (e.g., `neighboringElectrodesXXX.mat`) used for neighbor-based dissimilarity features. The appropriate file is selected automatically based on the number of channels in the input data. Currently the Module includes `.mat` files for EGI 128-channel and 256-channel nets; BioSemi 64-channel cap; and Easycap 74-channel cap.

#### ExampleData

By default, contains only a `readme.txt` file; due to file size, example datasets used by `example.m` are not included directly in the GitHub repository. Instead, `example.m` downloads the required files from SDR into this folder.

#### Figures

Default output folder used by `example.m` for saving generated figures (e.g., cluster visualizations, score plots, and UI screenshots).

#### HelperFunctions

Contains all helper functions called internally by `markSusChs()`. These functions implement windowing, feature computation, thresholding, clustering, and figure generation, and are not typically called directly by the user.

#### example.m

A complete walkthrough script showing how to configure the Module, run `markSusChs()`, and inspect outputs. This is the recommended starting point for new users.

#### markSusChs.m

The main user-called entry point. It computes feature-based suspiciousness scores, performs event-timing clustering, generates diagnostic visualizations, and launches the interactive review interface for final bad-channel decisions.

#### README.md

A brief overview of the Module, including expected inputs/outputs and pointers to documentation.

#### LICENSE

License for the repository (MIT).

#### User Manual.pdf

The User Manual for the current release, including background, setup instructions, a full walkthrough of `example.m`, interpretation guidance, and documentation of helper functions.

#### .gitignore

Git configuration file (no user action required). Notably, contents of ***ExampleData*** are ignored by design.

## Functionalities of the Module

The Bad-Channel Detection Module is a function-based MATLAB framework for EEG quality control that combines (i) multi-feature suspiciousness scoring, (ii) clustering based on co-occurrence of high-amplitude transient events, and (iii) interactive human validation. The Module is intended to be used after basic filtering and prior to spatial decomposition and multichannel analyses.

### Main entry point

In order to use the *Bad-Channel Detection Module*, the repository needs to be in the user’s MATLAB path.

#### Main Function

All computations and the interactive review are invoked through a single user-called function `markSusChs()`.

~~~
[susMask, badChList,] = markSusChs(…
X, fs, INFO, EOG, …);
~~~

#### Inputs

The primary input is an EEG matrix *X* ∈ ℝ^*C×T*^ (channels × samples) with sampling rate *f*_*s*_. If data have already been epoched into trials, trials should be concatenated along the sample dimension. A montage-specific neighbor map defines local neighborhoods 𝒩(*c*) f or channel *c* (used by neighbor-based features). Electrooculogram (EOG) channels must be specified as channel indices (or an `EOG` struct) to assist in identifying eye-related cluster during review. Functions also often involve numerous additional and/or optional inputs, which are detailed in function docstrings and in the User Manual.

#### Outputs

The function returns (i) a tri-state channel status vector, (ii) the final list of bad channels selected during interactive review, and (iii) a plotting payload struct with feature values, clustering outputs, and metadata. When figure saving is enabled, the Module exports key diagnostic visualizations (cluster views, feature-score plots, and review snapshots) using a consistent filename prefix.

We now describe the internal computational steps implemented within `markSusChs()`, focusing on how features are computed, combined, and used to automatically tag suspicious and bad channels (summarized on Figure 1). Detailed function-level specifications, i nputs/outputs, a nd u sage examples are provided in the User Manual hosted in the GitHub repository (see footnote 1).

### I. Multi-feature suspiciousness scoring

1. The Module computes complementary features intended to capture distinct channel failure modes. These features are used to prelabel channels as *good, suspicious*, or *bad*. Importantly, these labels are *suggestions* that are finalized during interactive review.

#### Robust per-channel standardization

For time-resolved operations, the signal is robustly standardized *within each channel over time* using the median and median absolute deviation (MAD):

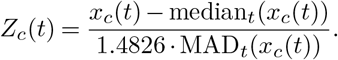

This produces a standardized signal *Z* that is used for detecting high-amplitude transient events for event-timing clustering, and computing windowed variance features on a common, channel-independent scale. This operation is implemented in helper function `zScoreRobust()` and stabilizes scaling in the presence of large transients.

#### Windowing and notation

Several features are computed over time windows. Let *w* ∈ {1, …, *W*} index windows of length *L* samples with hop size *H* samples. We denote the windowed segment for channel *c* by 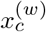. In the codebase, window formation is handled by helper function `makeWindowsFromX()`.

#### Feature 1. Neighbor dissimilarity (time-resolved)

For each window *w* and channel *c*, the Module computes Pearson correlations between *c* and each neighbor *j* ∈ 𝒩 (*c*). A robust neighbor similarity per window is then summarized as the median correlation across neighbors:

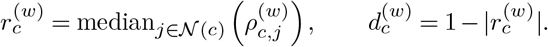

This yields windowed neighbor-dissimilarity values 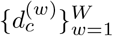.

#### Smoothing

The windowed series 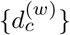 is smoothed across the window index using a moving-average operator (implemented in helper function `smoothFeature()`). Smoothing is used to reduce sensitivity to single-window fluctuations.

#### How it flags channels

After smoothing, the Module aggregates across windows to obtain a single per-channel neighbor-dissimilarity score, for example:

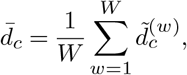

where 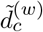 denotes the smoothed values. A channel is pre-flagged as suspicious if any window is greater than or equal to a predetermined threshold:

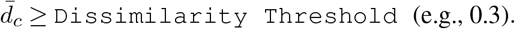

#### Feature 2. Extreme amplitude (hard failure) and elevated amplitude (soft warning)

Amplitude-based screening is split into a hard cutoff and a standardized warning score.

#### Hard cutoff (pre-label as bad)

Anecdotally, we have found that high-amplitude non-EOG transients degrade the performance of downstream ICA calculations. To account for this, for each channel *c*, the Module computes the maximum absolute amplitude:

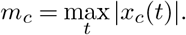

If *m*_*c*_ ≥ `bad-channel amplitude threshold` (e.g., 1000 µV), the channel is pre-labeled as *bad*. This criterion targets channels exhibiting extreme deviations from the typical signal range.

#### Standardized amplitude score

To detect channels with unusually large excursions that may not cross the hard cutoff, the Module computes a standardized score across channels using a log transform and z-scoring:

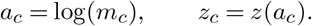

If *z*_*c*_ ≥2 and not marked as *bad*, the channel is pre-labeled as *suspicious*.

#### Feature 3. Overall Variability and Temporal Variability

Variance-based screening targets channels that are unusually noisy, unusually flat, or unstable over time.

#### Global variance level (time-aggregated)

The Module computes per-channel variance *v*_*c*_ = Var(*x*_*c*_), applies a log transform, and robustly *z*-scores across channels:

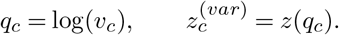

High variance indicates noisy or unstable electrodes, while very low variance may reflect flat or poorly conductive channels. Both situations are undesirable. The Module therefore, keeps both tails of the distribution, but uses a higher magnitude theshold on the lower side (-2.5) because low-variance channels can occur naturally in quieter scalp regions or near the reference; they should only be flagged when they are clear outliers relative to the group:

If 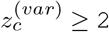 or 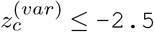, channel *c* is pre-labeled as *suspicious*.

#### Temporal Variability

Variance is also computed within windows, producing 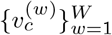. After smoothing across windows (again via `smoothFeature()`), the Module summarizes instability using a range statistic:

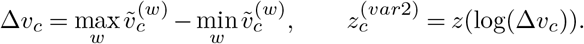

If 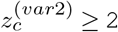, channel *c* is pre-labeled as *suspicious*.

### II. Clustering based on co-occurrence of high-amplitude transient events

A central goal of the Module is to *expose shared structure* among channels that exhibit temporally aligned, high-amplitude transient events. To do so, the Module converts each channel into a sparse event-timing representation, computes pairwise channel similarity in that representation, and clusters channels by thresholding the resulting distance matrix. This clustering is used for *organization and interpretability* during review, rather than as an automatic classification rule (see User Manual for parameter defaults and additional examples).

#### (1) Robust z-scoring and high-amplitude transient detection

The procedure begins by robustly standardizing each channel using helper function `zScoreRobust()`. This step is essential to prevent high-amplitude transients from inflating the standard deviation.

A binary transient mask is then formed by thresholding the magnitude of the robust *z*-score:

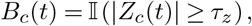

where *τ*_*z*_ is a user-configurable threshold (e.g. *τ*_*z*_ = 14). Then, *B*_*c*_(*t*) = 1 denotes time points where channel *c* exhibits unusually large deviation relative to its typical within-channel variability.

#### (2) Temporal max-pooling to obtain an event-timing vector

To represent transient timing at a coarser temporal resolution and reduce redundancy, the Module performs max-pooling over short, non-overlapping blocks of duration *l* seconds (e.g *200ms*). Let ∈ *b* {1, …, *B*} index pooled blocks; then the pooled binary event indicator is:

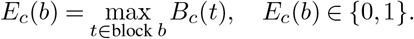

This step converts each channel into a sparse event-timing vector *E*_*c*_ ∈ {0, 1}^*B*^.

#### (3) Pairwise channel distance from pooled event masks (overlap coefficient)

After robust thresholding and maxpooling, each channel *c* is represented by a binary vector *b*_*c*_ ∈ {0, 1}^*B*^, where *b*_*c,k*_ = 1 indicates that channel *c* contains at least one suprathreshold transient within pooled block *k*. Let

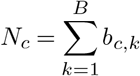

denote the number of transient-active blocks for channel *c*. For a pair of channels (*c*1, *c*2), let the intersection count be

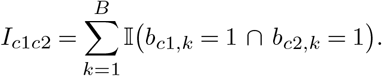

We define a Jaccard-like (overlap-based) distance

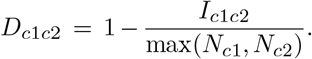

If max(*N*_*c*1_, *N*_*c*2_) = 0 (neither channel exhibits any detected transients), we set *D*_*c*1*c*2_ = 0 so that transient-free channels are treated as maximally similar. This computation is implemented in helper function `jaccardDistanceChannels()`.

#### (4) Clustering via distance thresholding

Given the distance matrix *D*, the Module forms an adjacency graph by thresholding distances at *ε*:

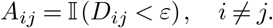

Clusters are then defined as the connected components of this graph. In other words, channels belong to the same cluster if they can be linked through one or more edges connecting sufficiently similar event-timing patterns. This step is implemented in helper function `clusterByThreshold()` (see Figure 2).

**Fig. 2.**
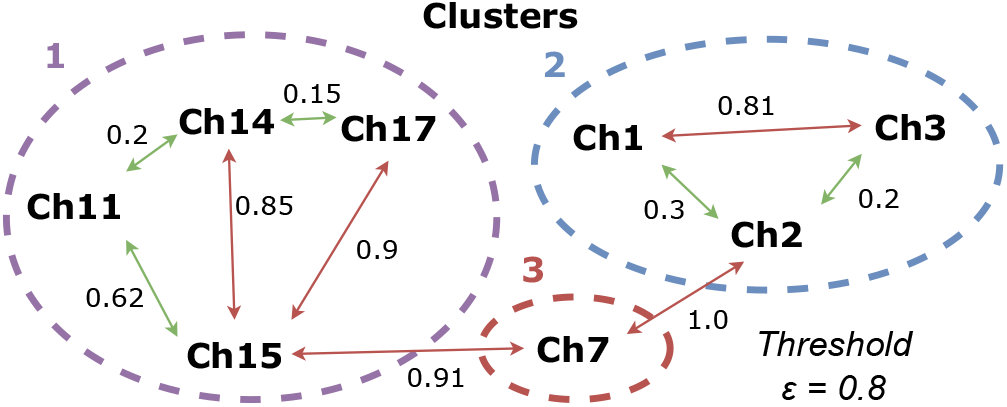
Clustering via distance threshold. Edges connect channel pairs whose high-amplitude transient pattern distance is below the threshold *ε*. Clusters correspond to the connected components of this graph.

**Fig. 3.**
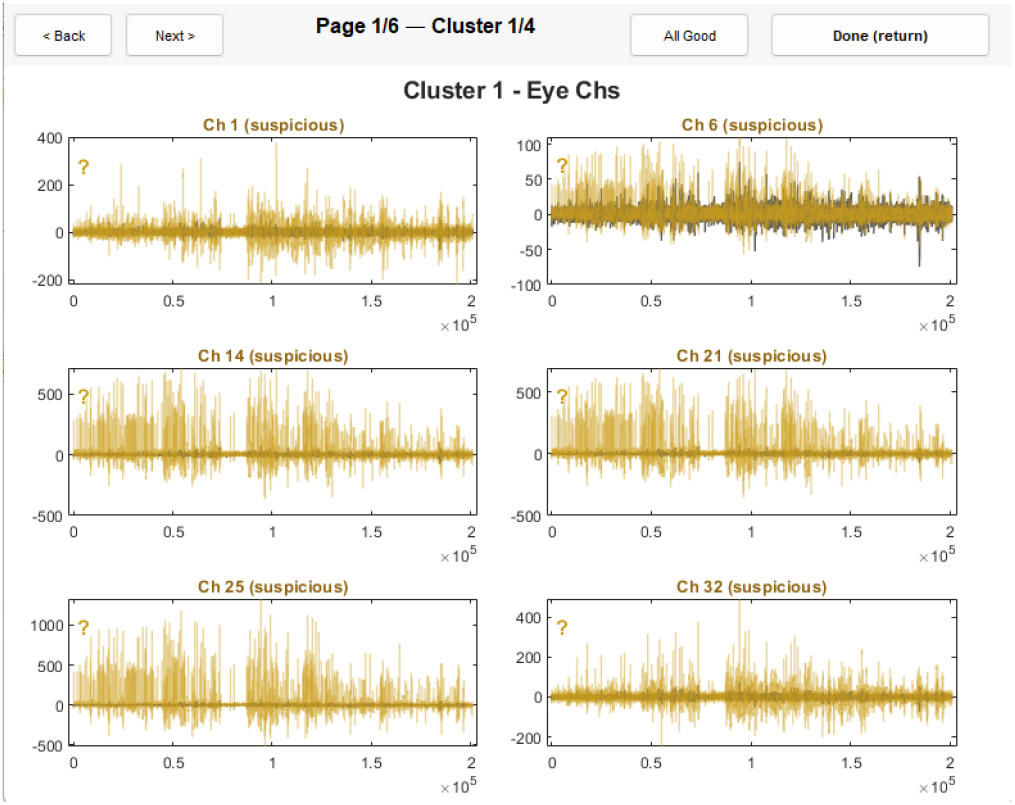
Interactive review interface (example). Channels are organized to support rapid validation of pre-flagged channels and cluster-level patterns. Final classification remains user-defined. See the User Manual for interface controls (button functions and shortcuts), cluster organization details, and practical guidance for interpreting traces and deciding whether a channel should be marked good, suspicious, or bad.

#### Interpretation

These clusters reflect *shared transient-event timing*, which may arise from common artifact sources or from structured physiological signals. Clusters are therefore treated as an organizational tool for review rather than an automatic decision: clusters may contain good channels, bad channels, or mixtures depending on context.

### Eye-related grouping

The Module identifies the primary cluster that overlaps with EOG indices as the *eye-related* group and constrains its pre-labeling (i.e., prevents eye cluster from being auto-labeled as *bad* based purely on amplitude). This is designed to reduce conflating *bad*-channel categorizations with physiologically meaningful ocular structure while still presenting these channels prominently for review.

### Small Clusters

In particular, clusters containing a small number of channels are preferentially selected for manual inspection, as small clusters are more likely to reflect channel-specific failures rather than spatially distributed artifacts.

### III. Interactive review and final bad-channel list

Automated feature screening and transient-event clustering are used to *prioritize* and *organize* channels for inspection, but the Module does not enforce fully automatic channel rejection. Instead, it concludes with a human-in-the-loop validation stage implemented by the interactive GUI `reviewBadChsUI()`, which is launched from within `markSusChs()`. The goal of this step is to reduce the burden and subjectivity of manual channel inspection by (i) prioritizing channels flagged by multiple features and (ii) grouping channels that share transient-event timing, so users do not need to manually scan recordings to locate similar events across channels. The GUI further streamlines decision making by enabling interactive finalization of the bad-channel list, avoiding ad hoc script edits (e.g., manually hard-coding channel indices).

#### Tri-state labeling and decision logging

The interface supports tri-state labels:

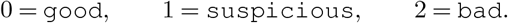

The initial state is populated from automated feature screening, and the user can interactively toggle channel states while inspecting time-domain overlays. The *suspicious* state denotes “needs attention” but does not automatically flag the channel for removal; the user must manually toggle such channels to *bad* to confirm removal. The interface records the final label for every channel, enabling a complete audit trail of what was inspected and how channels were classified.

#### Final output rule

The final bad-channel list returned by `markSusChs()` includes only channels explicitly marked as *bad* (state 2) at the conclusion of interactive review:

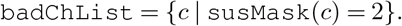

## Illustrative analysis

This illustrative analysis demonstrates the user-facing workflow of the Bad-Channel Detection Module: obtaining example data, running the main entry point, and interpreting the figures and interactive review interface that lead to the final bad-channel list. The intent is to show what users will see and how the outputs should be read at a high level; detailed button descriptions, UI controls, and channel-labeling guidelines are provided in the User Manual.

### Running the Module

To ensure the analysis is fully reproducible, the GitHub repository includes a runnable script, ***example*.*m***, which calls the Module’s main entry point, `markSusChs()`. Users can run the illustrative analysis without modifying code paths beyond setting the repository location and an output directory for figures (if saving is enabled).

### Example data and how it is obtained

The example script automatically downloads the example EEG recording(s) from the Stanford Digital Repository (SDR) into the local ***ExampleData*** folder. The example data represent adult data recorded from the Electrical Geodesics, Inc. (EGI) GES 300 platform [6]. The script then loads the continuous EEG matrix and accompanying metadata (sampling rate and EOG channel specification) required by the Module.

### What the user sees: figures and interface

When the Module is run, the user is presented with a consistent set of visual outputs designed to support rapid and defensible channel decisions. The figures are intended to be read in the order below.

#### (1) Cluster visualization: “Which channels share the same transient pattern?”

The first figure (Figure 4) visualizes the channel clustering derived from shared transient-event timing. It includes (i) a graph-based cluster view and (ii) a distance matrix/heatmap reordered by cluster membership. Both panels depict the clusters structure in complementary forms. The largest cluster (C1) appears as the largest group in the graph-based view and as a uniformly blue square in the reordered distance matrix. The near-zero distances within this region indicate that channels within C1 have identical high-amplitude transient masks; in practice, this most often reflects the absence of such transients across those channels. Beyond showing the resulting groups, this figure serves as a diagnostic of clustering quality: it provides immediate visual feedback on whether clusters are well separated under the current parameterization. The User Manual describes recommended tuning strategies and the relevant parameters.

**Fig. 4.**
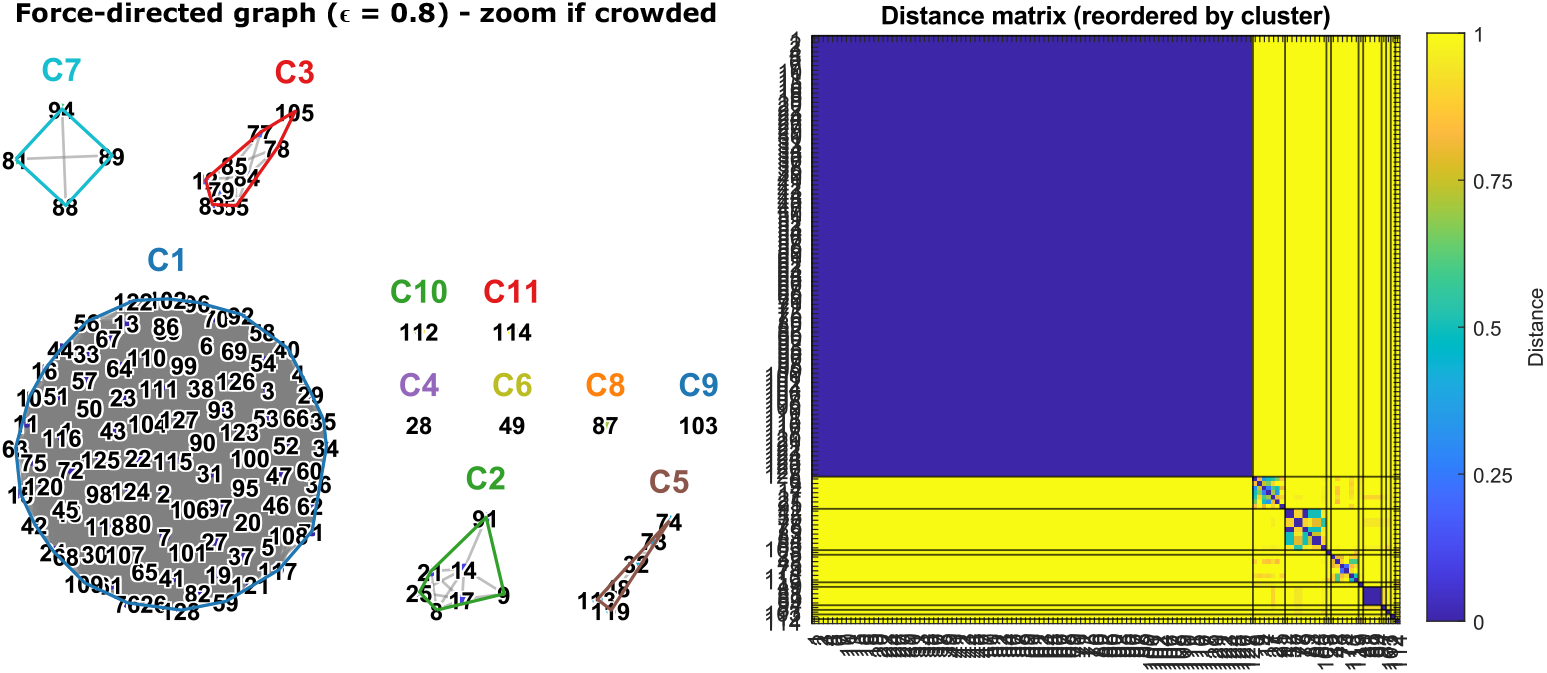
High-amplitude transient pattern clusters and distance matrix. *Left:* Force-directed graph showing high-amplitude transient pattern clusters. *Right:* Distance matrix reordered by cluster index. Cluster C1 is represented by the large blue square on right plot of. Channels in this cluster are less likely to be bad or suspicious. Clusters C-10, 11, 4, 6, 8, and 9 are singletons (one-channel clusters). Clusters C-7, 3, 2, and 5 are small channel clusters tagged for review. Default: if number of channels per cluster is less than 7, then the whole cluster is tagged for review.

#### (2) Feature-score summary: “Which channels should I look at first?”

The second figure (Figure 5) summarizes the feature-based pre-flagging stage. It visualizes per-channel scores from the screening features (e.g., neighbor dissimilarity, amplitude checks, and variability measures) and highlights channels that are most likely to warrant inspection. The goal of this plot is *triage*: it narrows attention to a small subset of channels rather than requiring uniform manual scanning across the entire montage.

**Fig. 5.**
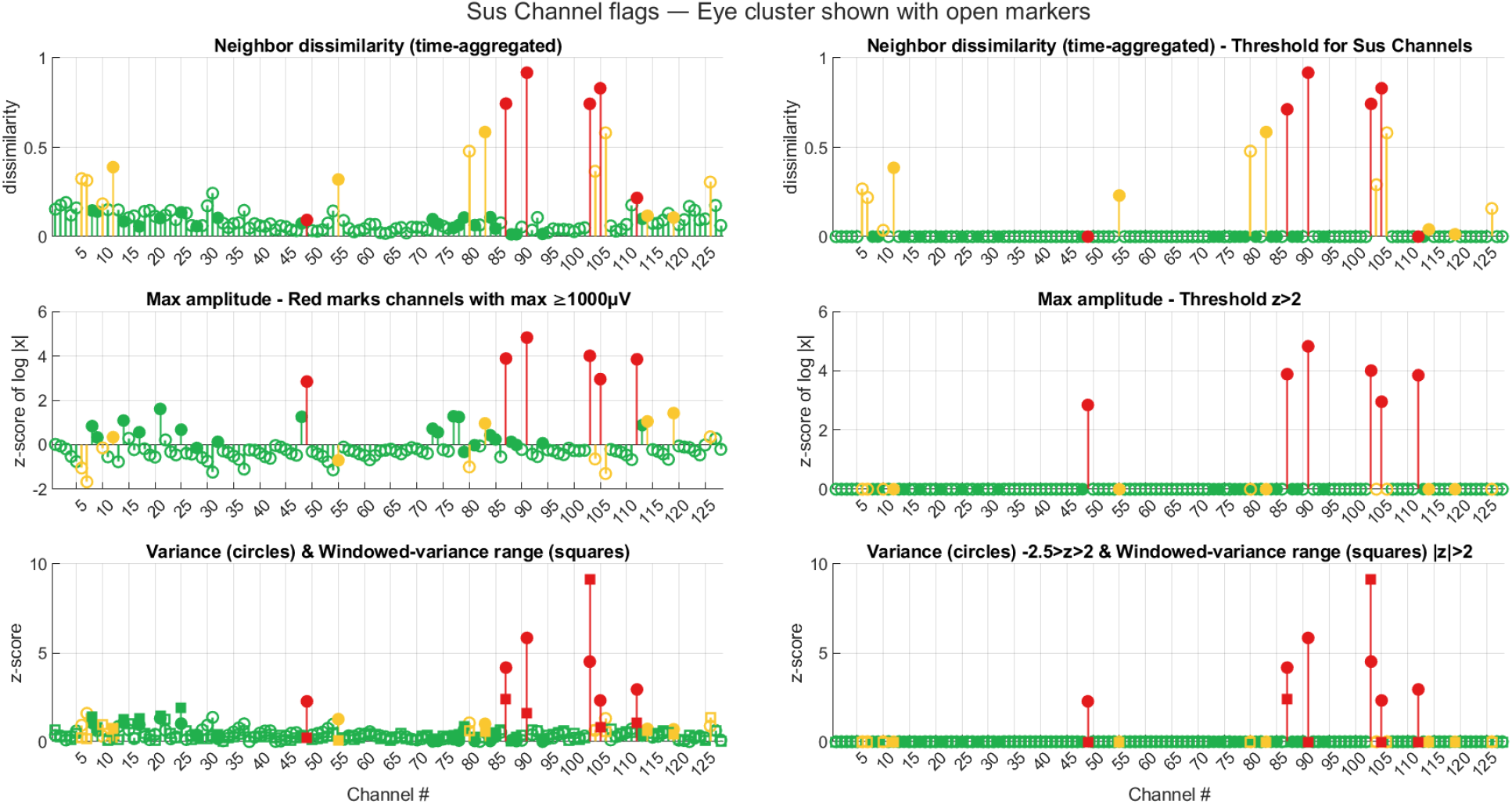
Feature-score summary. *Left*: Feature scores per channel. *Right*: Channels exceeding suspicious /bad thresholds. There are multiple Suspicious and Bad channels, and this recording needs detailed inspection. Remember to reference these flags when assessing a channel and deciding whether to tag it as bad. See the User Manual for feature definitions and threshold behavior.

#### How to interpret it (high level)

Channels that appear as clear outliers across one or more feature panels are strong candidates for review. Channels that are consistently non-outlying are typically deprioritized. This plot is not a final decision rule; it is a prioritization view (see User Manual for feature definitions and threshold behavior).

#### (3) Interactive review UI: “Finalize the bad-channel list”

The UI presents channels in an order designed to accelerate review: cluster-guided, with suspicious candidates emphasized. Users inspect time-domain traces and toggle channel status (*good /suspicious /bad*). The review step concludes when the user selects the UI control to finalize and return outputs to MATLAB.

#### Output rule

Only channels explicitly labeled as *bad* at the end of interactive review are included in `badChList`. Channels left as *good* or *suspicious* are retained.

#### UI Panels and Manual Decisions

For readability, the main text presents only the representative UI overview in Figure 6. A complete panel-by-panel walkthrough of this illustrative example, including cluster-specific decisions and before/after views, is provided in the Supplementary Information.

**Fig. 6.**
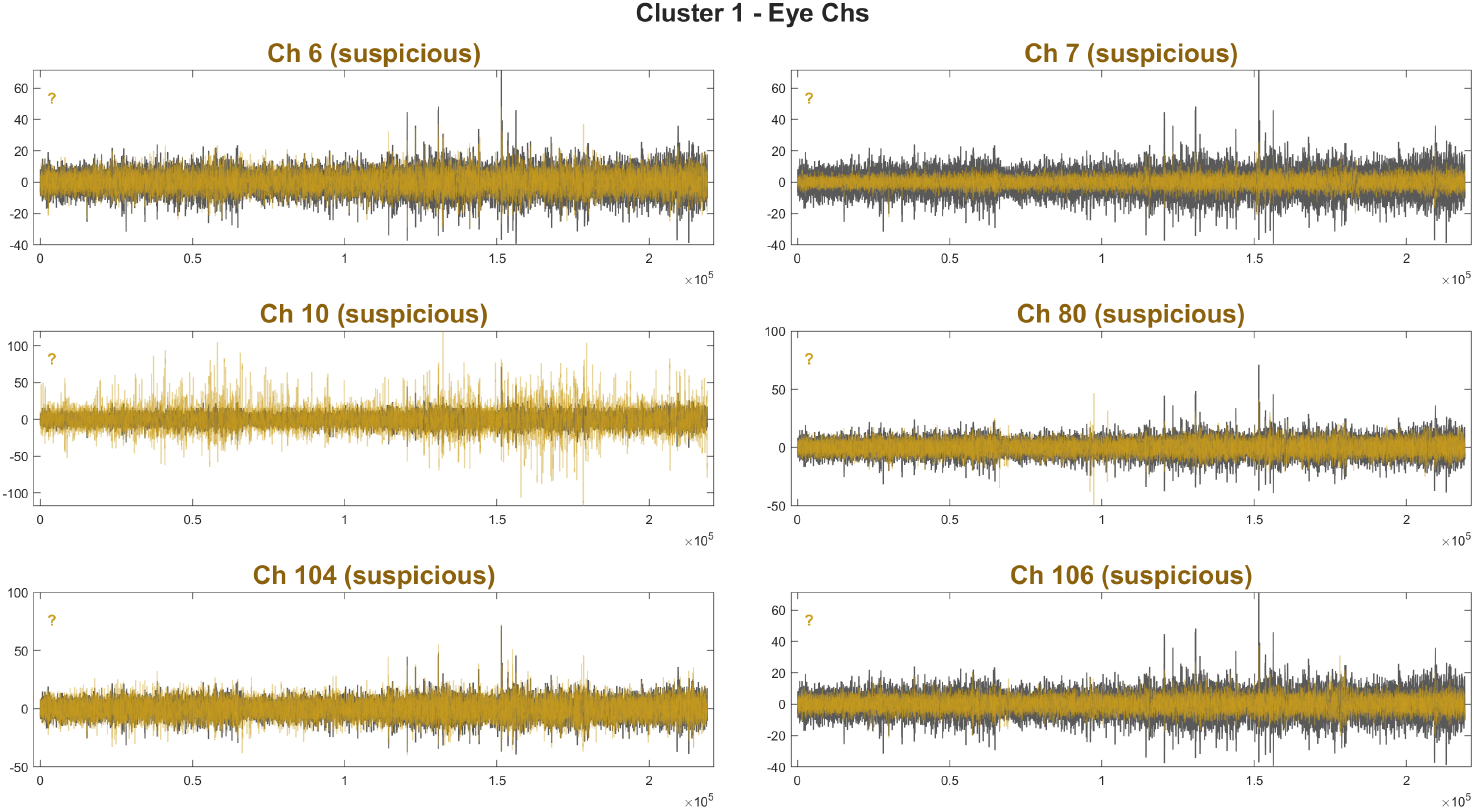
Interactive review interface (representative view). Channels are organized to support rapid validation of pre-flagged channels and cluster-level patterns. Final classification remains user-defined. For descriptions of UI controls (buttons/shortcuts), grouping/ordering behavior, and recommended criteria for labeling channels, refer to the User Manual.

**Fig. 7.**
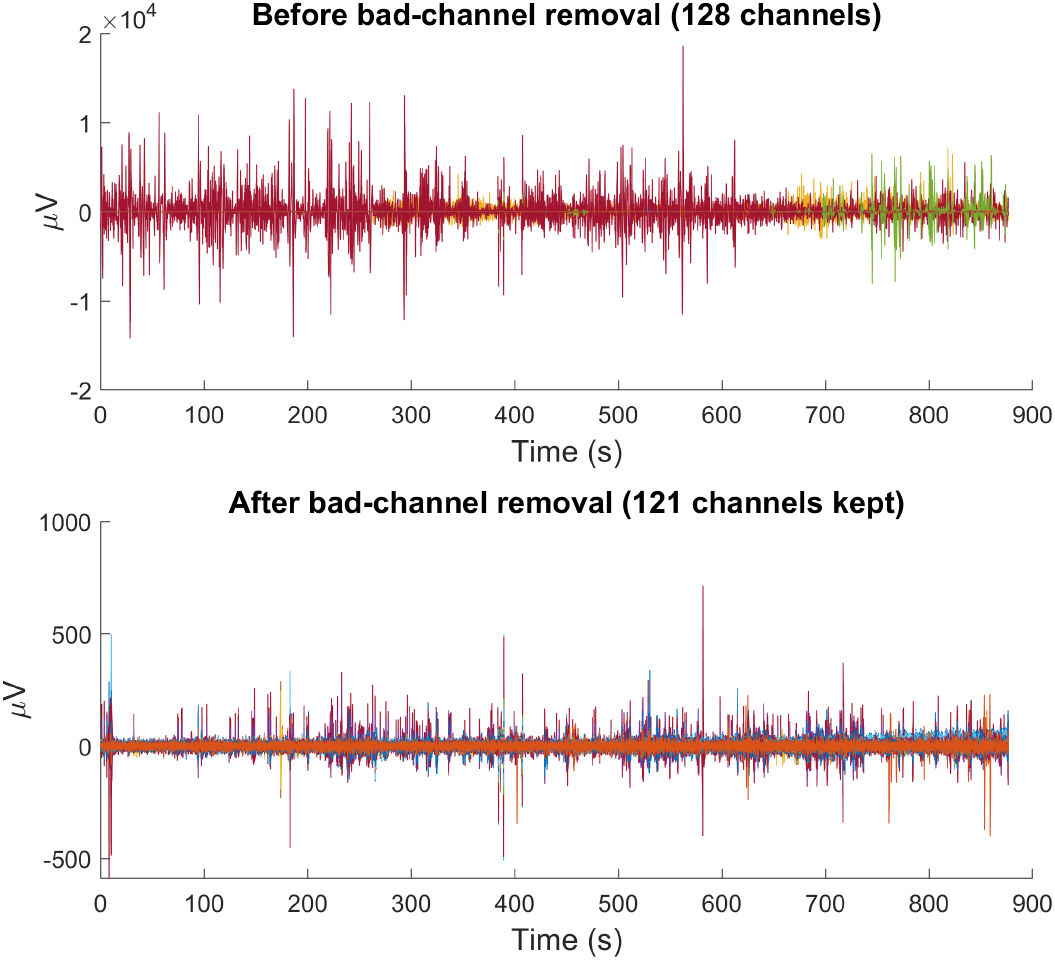
Before/after summaries. Time-domain traces from all channels are overlaid in a single axis. In this example, the recording contains 128 channels and 7 were marked as bad during interactive review. Removing these channels substantially reduces the amplitude scale required to display the overlay: the y-axis range changes from approximately ±20,000 *µ*V (before) to ±1,000 *µ*V (after), reflecting the removal of extreme excursions that would otherwise dominate summary views. This figure is intended as a qualitative sanity check and for documentation, not as a standalone decision rule.

#### (4) Optional before/after summaries: “What changed after removal?”

To support quick sanity checks, the Module can generate summary plots comparing the recording before versus after removing user-marked bad channels. These plots are not meant to prove correctness; rather, they provide a rapid qualitative check that obvious failures (e.g., extreme excursions or unstable channels) are no longer dominating the recording summaries.

Further elaboration and interpretations for this analysis are provided in the User Manual.

## Discussion

We presented the first release of the *Bad-Channel Detection Module*, a MATLAB codebase for EEG quality control that combines (i) multi-feature screening, (ii) clustering based on shared transient-event timing, and (iii) an interactive review interface that preserves user control over final channel removal. Bad-channel handling is widely recognized as a critical step in EEG preprocessing because a small number of malfunctioning sensors can dominate time-domain visualizations, inflate amplitude scales, and propagate contamination into downstream stages of data cleaning and analysis [3].

A central design decision of the proposed Module is that channels exhibiting similar high-amplitude transient structure should not automatically classified as bad. While isolated transients or persistent deviations localized to a sin-gle channel are strong indicators of electrode failure, transient events that co-occur across channels are more consistent with global or spatially distributed artifacts. These patterns are precisely those targeted by component-based cleaning approaches such as ICA, which benefit from preserving shared variance across electrodes. By clustering channels based on the temporal overlap of extreme events and routing these clusters to human review rather than automatic rejection, the proposed approach avoids conflating shared artifact structure with channel failure and preserves information critical for downstream decomposition.

### Relation to existing approaches

The present Module is not the first approach to EEG bad-channel identification, and users have many options depending on workflow goals and desired level of automation. Several established toolboxes and pipelines implement multi-feature channel screening and, importantly, many go beyond a single global channel statistic by incorporating time-local evaluations. For example, FASTER applies a fully automated cascade of outlier screens at multiple levels (including channel-level statistics and epoch-wise evaluations), using standardized feature scores to flag channels/segments/components for rejection or interpolation [2]. PREP similarly combines multiple complementary channel-quality criteria and includes a time-windowed predictability test based on RANSAC-style channel reconstruction, which can identify channels that become poorly explained by subsets of other channels during portions of a recording [3]. The EEGLAB ecosystem provides automated detection of problematic channels (e.g., flatline/noisy channels) and is often paired with Artifact Subspace Reconstruction (ASR), which aims to attenuate high-variance artifact subspaces relative to a reference “clean” data segment [1].

Despite their utility, fully automated routines can still underperform in realistic laboratory data regimes for several practical reasons. First, the distinction between a truly malfunctioning sensor and a channel containing recoverable contamination can be ambiguous: some extreme or intermittent events may later be sufficiently handled by downstream steps (e.g., ICA), whereas other events are large enough to dom-inate preprocessing and warrant channel removal. Second, time-varying failures can be difficult to separate cleanly from typical variability using a single set of thresholds across heterogeneous recordings, leading to both missed bad channels and overly aggressive rejection depending on parameterization and artifact prevalence. Third, even when a pipeline computes time-local statistics, its decision rule often produces a binary outcome (keep vs. reject/interpolate) without a convenient workflow for rapid human validation of borderline cases. As a result, many users still perform manual inspection even when automated methods are available, particularly for high-density montages and long recordings. The Bad-Channel Detection Module targets this practical gap by explicitly structuring the human validation step rather than attempting to eliminate it. The Module uses conservative screening thresholds so that channels with strong evidence of failure are at least pre-flagged as *suspicious* — reducing the chance they are silently missed.

For clustering bad channels based on the co-occurrence of high-amplitude transient events, this Module employs a custom derivative of the Jaccard similarity equation [7]. Jaccard-based similarity measures have been previously introduced in EEG analysis as a means of comparing discrete or symbolic signal representations, for example to quantify similarity between ordinal patterns [8], or to cluster channels based on shared brain rhythm sequences [9], demonstrating advantages over Pearson correlation when the goal is to characterize state-based behavior rather than linear amplitude relationships.

In contrast to these prior applications, which typically compute similarity over the full recording, our approach uses robust *z*-score thresholding (|*Z*| *>* 14) to generate binary high-amplitude transient masks that intentionally strip away most physiological activity, leaving a sparse representation of extreme, non-physiological events. Custom Jaccard similarity is then applied exclusively to the co-occurrence of these transients, rather than to extended periods in which both channels are quiet. This distinction is critical in high-density EEG, where bad channels often share intermittent, high-amplitude artifacts driven by common external sources such as localized motion, cable disturbances, or hardware instability. By ignoring jointly quiet intervals and emphasizing the intersection of transient events, the resulting similarity matrix preferentially identifies channels coupled by shared artifactual sources rather than by neural activity. This relational perspective contrasts with traditional bad-channel detection methods [1, 2] — which primarily evaluate channels in isolation using amplitude, variance, or correlation thresholds — and enables the identification of clusters of channels that are not globally corrupted but are intermittently affected by the same non-biological disturbance, supporting a more targeted and interpretable rejection strategy.

### Contributions and recommended usage

The primary contribution of this work is a streamlined, interpretable workflow for bad-channel quality control that makes channel decisions both faster and easier to justify. The Module combines complementary feature-based screening with a novel clustering method driven by the co-occurrence of robustly detected high-amplitude transients, allowing channels affected by shared artifact sources to be reviewed together rather than in isolation. This organization reduces the need to manually search for comparable events across channels and provides immediate diagnostic feedback on whether co-artifactual structure is well separated under the current parameterization. An interactive graphical interface further accelerates the review process by enabling rapid tri-state labeling (*good/suspicious/bad*) without modifying scripts or hard-coding channel indices, ensuring that final bad-channel lists reflect explicit user decisions while remaining efficient and well documented. Importantly, the Module returns — and can optionally save — all information used during review, including feature summaries, cluster assignments, and standardized visualizations before and after channel removal. Together, these design choices support transparent decisionmaking, reproducibility, and reportable quality-control outcomes, positioning the Module as a practical human-in-the-loop alternative to fully automated bad-channel rejection strategies.

The Module is not intended to replace expert judgment or serve as a fully automatic classifier. First, the clustering stage is most informative when artifacts manifest as intermittent transient events; datasets dominated by non-transient contamination (e.g., strong muscle activity, slow drifts, or broadly distributed noise) may yield weak cluster separation or clusters that reflect broad dynamics rather than sensor failure. Second, parameter choices can affect sensitivity and specificity; users should treat the clustering visualization as diagnostic when adapting the Module to new recordings and adjust parameters when clusters appear diffuse or implausible. Third, physiological sources (notably ocular activity) can produce large and structured signals that resemble artifacts under amplitude-based checks; the Module therefore emphasizes interactive validation and supports EOG-aware handling rather than automatic rejection of eye channels. Finally, neighbor-based measures assume a correct montage definition and indexing; errors in channel layout specification can degrade neighbor metrics and downstream interpretations. In practice, we recommend using the Module as an explicit quality-control step early in preprocessing to identify and remove clearly malfunctioning sensors and to document channel-level QC decisions. The *suspicious* label should be understood as a conservative pre-flag intended to focus review; channels are removed only when explicitly labeled as *bad* during the interactive step.

### Limitations and future work

We recognize limitations of the current release of the Module. First, it is important to note that this Module was developed as part of a larger data-cleaning pipeline designed to preprocess long EEG trials (on the order of minutes) in experimental paradigms where few to single trials are recorded from each participant. In these contexts, rejecting trials can be costly, and bad-channel identification helps to maximize the number of usable trials. However, users working with shorter epochs and/or higher trial counts may have more leeway to reject trials that are contaminated by artifacts. Moreover, users may not need the high-amplitude transient criterion if such transients do not impact their own downstream preprocessing steps, or if the user can interpolate small time segments rather than reject the channel altogether. The workflow includes an interactive review stage requiring user intervention; this may not be preferred by users seeking full automation. Finally, `neighboringElectrodesXXX.mat` files are available only for selected montages (EGI 128 and 256 channel and derivatives; BioSemi 64-channel; and Easycap 74-channel [10]); users of other montages will need to create their own `.mat` files in order to compute neighbor-based dissimilarities.

Future improvements may focus on making clustering behavior more robust across recordings without weakening the conservative screening philosophy. First, the clustering distance threshold (and related pooling settings) could be selected more automatically based on properties of the distance matrix or event-rate statistics, reducing user tuning burden. Second, cluster summaries could explicitly highlight within-cluster outliers (channels that cluster with an artifact family but remain atypical relative to the cluster majority) to further accelerate review. Third, the current binarization step (following per-channel robust standardization) can yield slightly different event masks for channels that are visually similar but have different baseline dynamics; improving this mapping (e.g., adaptive binarization tied to per-channel dynamics) may reduce unintended cluster splitting and improve cluster interpretability. These changes would strengthen the Module’s primary goal: rapid, defensible channel decisions supported by clear, reviewable evidence.

## ACKNOWLEDGEMENTS

This work supports the Open Science efforts of the Stanford Educational Neuroscience Initiative (SENSI). The authors thank members of SENSI and collaborating labs at Stanford University for their feedback and early testing of the Module. We especially acknowledge Philip Hernandez, Neha Rajagopalan, and Madison Bunderson for their insights delivered from having performed manual review and classification of bad channels in SENSI’s EEG datasets. Philip Hernandez additionally provided detailed feedback on early algorithm behavior, including the rationale for channel rejection and identification of failure modes that informed subsequent refinements.

This preprint was prepared using the *HenriquesLab bioRxiv* Overleaf template by Ricardo Henriques.^6^

## AUTHOR CONTRIBUTIONS

Designed the algorithm and implemented the code: AJM. Validated and integrated into larger *SENSI EEG PREPROC* pipelines: AJM, BK. Documented the Module: AJM. Created illustrative analyses: AJM. Wrote the preprint: AJM. Provided supervision, feedback, and editing: BK.

## DECLARATION ON THE USAGE OF AI

The authors used OpenAI’s GPT-5 and Google’s gemini-2.5-flash models for assistance with documentation formatting, function summaries, LaTeX structure, and for brainstorming and exploring conceptual ideas. Additionally, these tools were used to improve flow, word choice, and structure of the User Manual and preprint. No AI tools were used for data analysis, algorithm development, evaluation, or initial drafts of the User Manual or preprint.

## Supplementary Note 1: Illustrative analysis — Supplementary UI panels

**Fig. S1.**
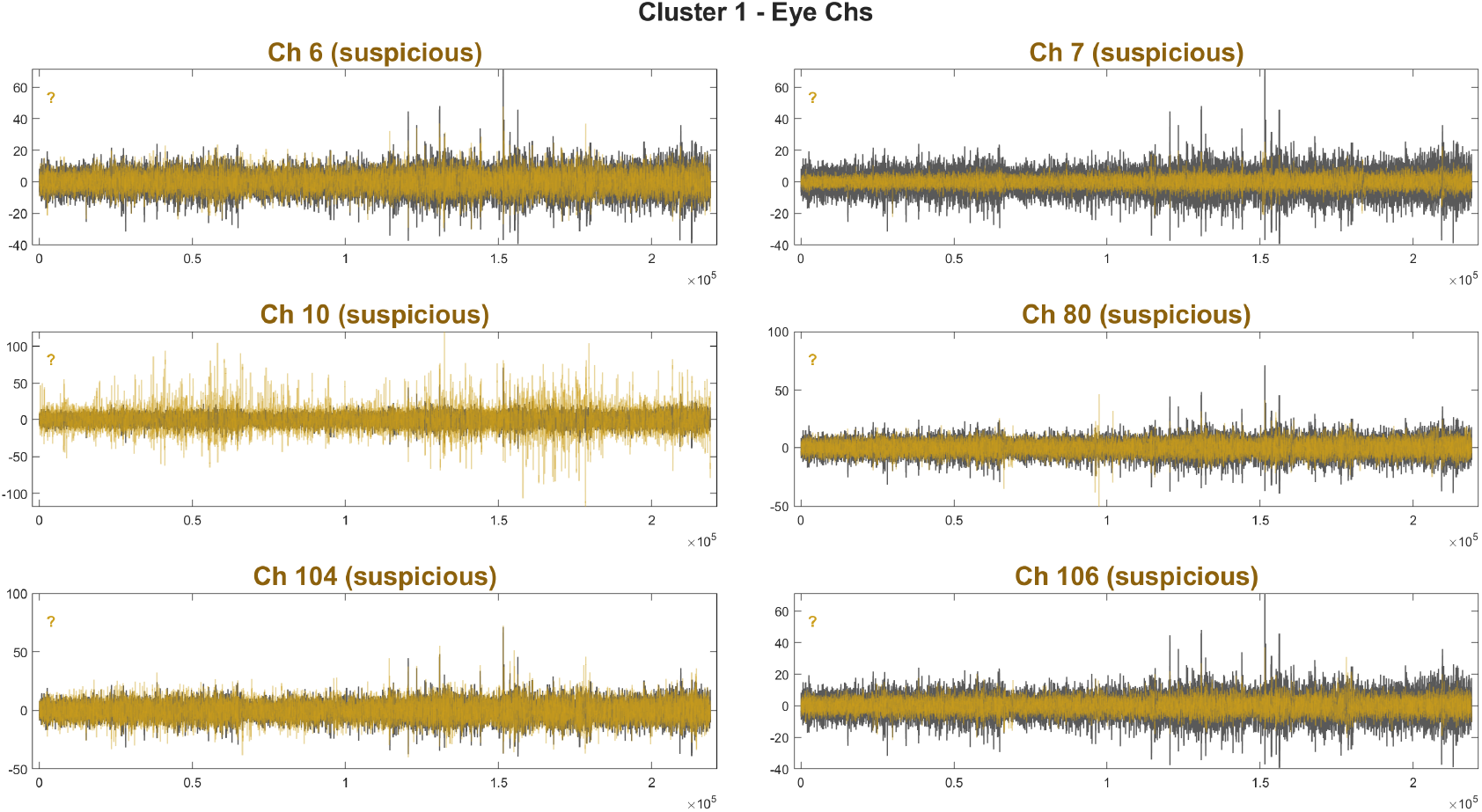
Page 1, cluster 1 — eye channels. We consider these channels to be acceptable and would not flag any for rejection. Channel 10 is slightly noisier than the rest, but not an atypical waveform for an EOG channel and lacks flags in Figure 5. Channel 106 had suspicious Neighbor dissimilarity (>0.4), but the waveform looks acceptable.

**Fig. S2.**
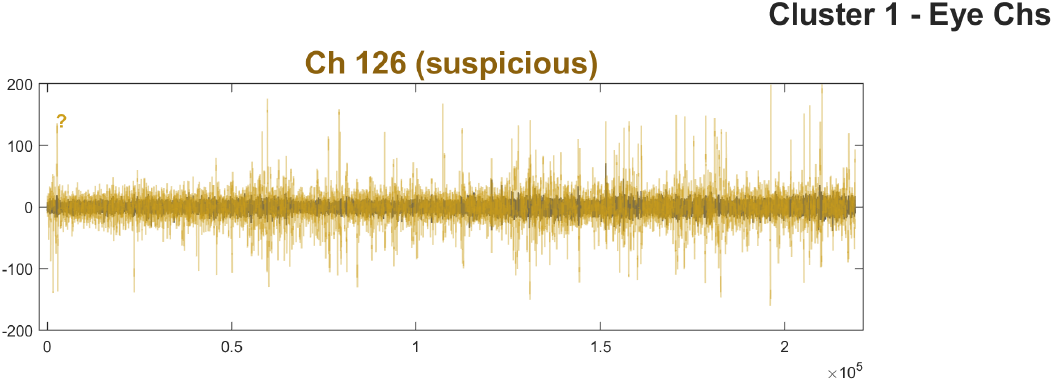
Page 2, cluster 1 — eye channels. We also consider this channel to be acceptable. It is similar to channel 10 but with higher amplitude; placed on the cheek, this channel is sometimes used for VEOG. It has no strong flags.

**Fig. S3.**
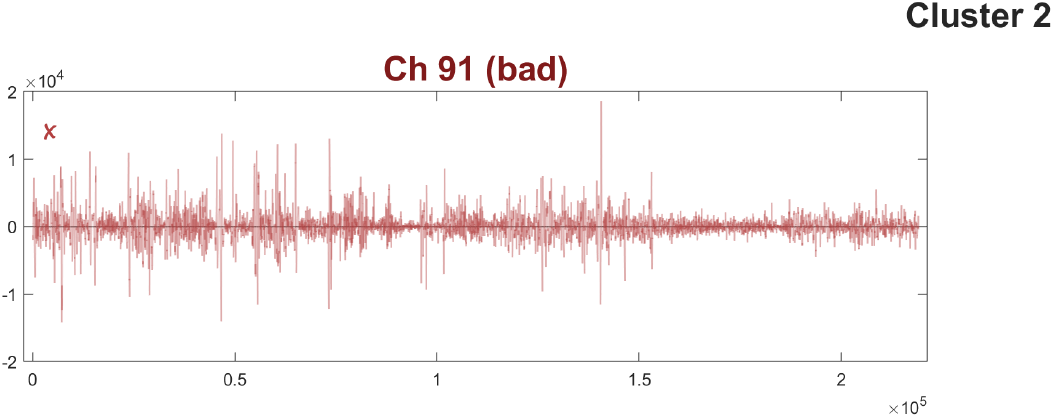
Page 3, cluster 2 (singleton). This channel was successfully auto-tagged as *bad*. The shown data are not characteristic of EEG activity but rather reflect a high-amplitude artifact (the electrode may be broken or have poor connection with the scalp.) This channel had the highest neighbor dissimilarity in Figure 5.

**Fig. S4.**
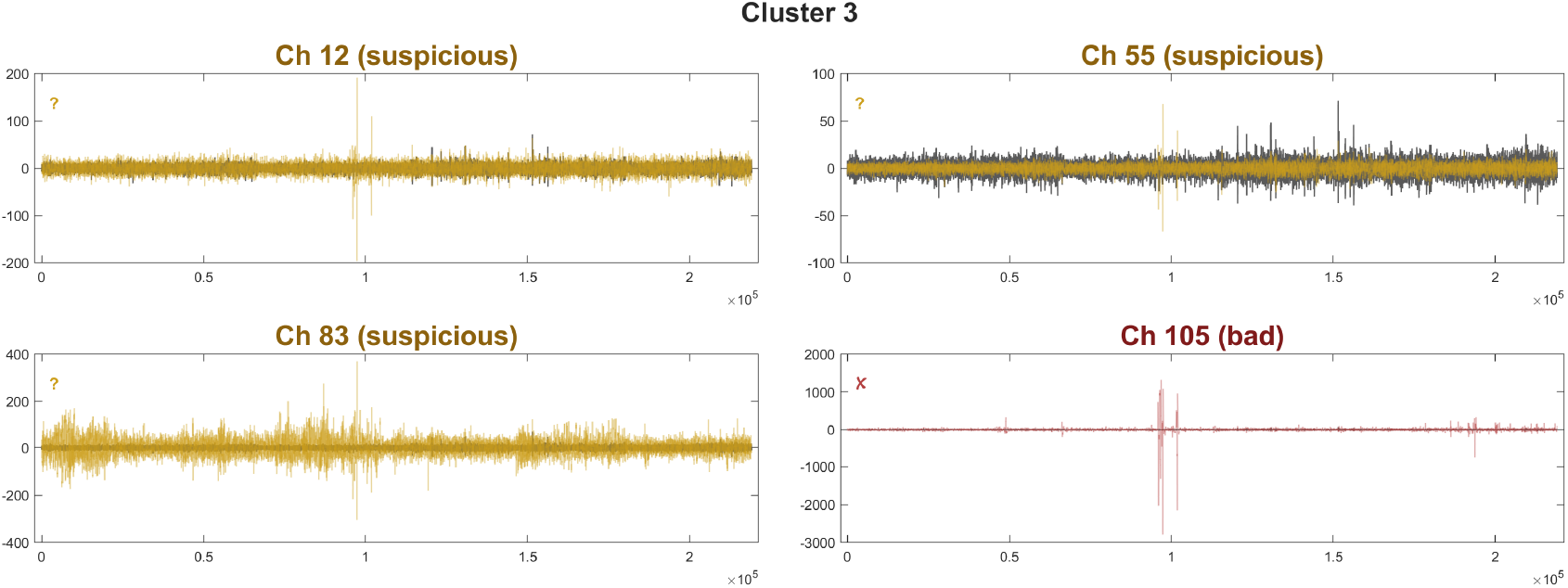
Page 4, cluster 3. We evaluate channels 12 and 55 as acceptable. Channel 83 is slightly noisier, with a high neighbor dissimilarity score of ≈ 0.71. Due to this, we will flag this channel for removal by clicking on the plot until it toggles to red. Channel 105 has two transients located in the same position as other channels in this cluster, and it is most likely the source of this cluster’s high-amplitude transients. However, the magnitude is too large and in our broader workflow, it would affect ICA performance. Therefore, we will remove channel 105 as originally tagged.

**Fig. S5.**
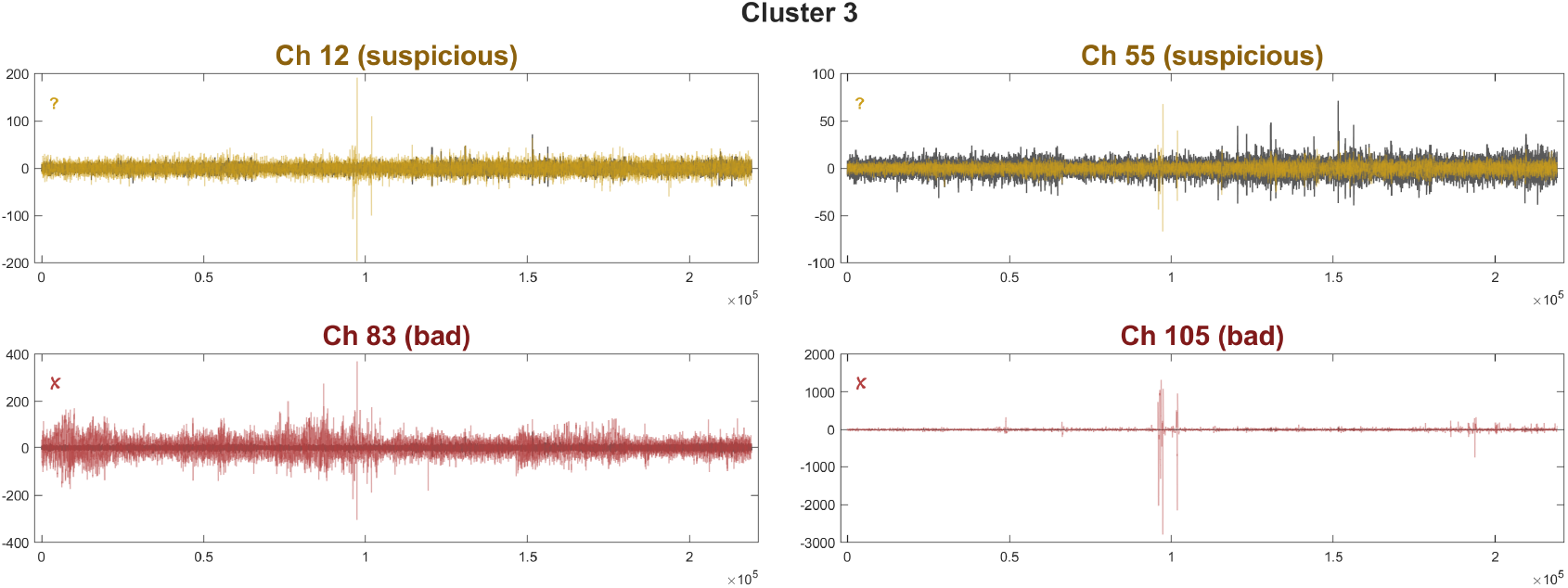
Page 4, cluster 3 (after review). This is how page 4 (formerly Figure S4) should look after user review and intervention. Channel 105 was already tagged as *bad*, and we added channel 83 for removal.

**Fig. S6.**
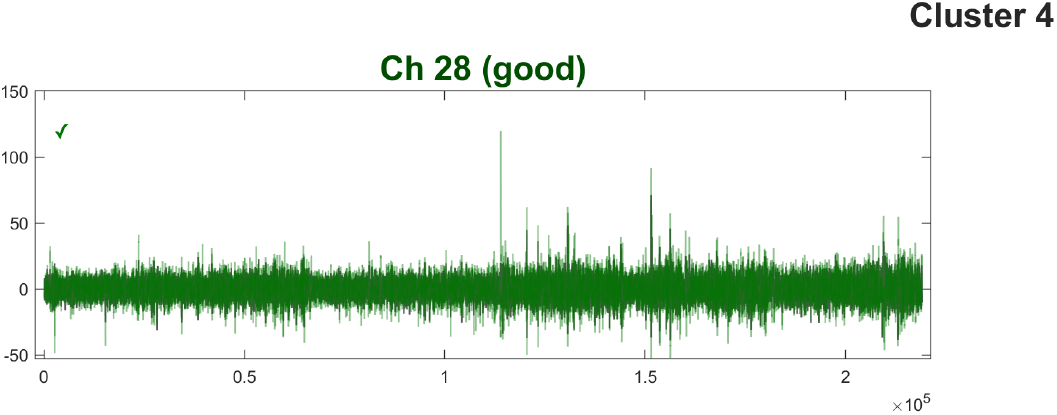
Page 5, cluster 4 (singleton). We consider this channel to be acceptable. It has “small” transients that also appear in other channels, and voltages are low enough that they will likely not impact ICA. It is likely that these artifacts can be removed by other means without needing to reject the entire channel.

**Fig. S7.**
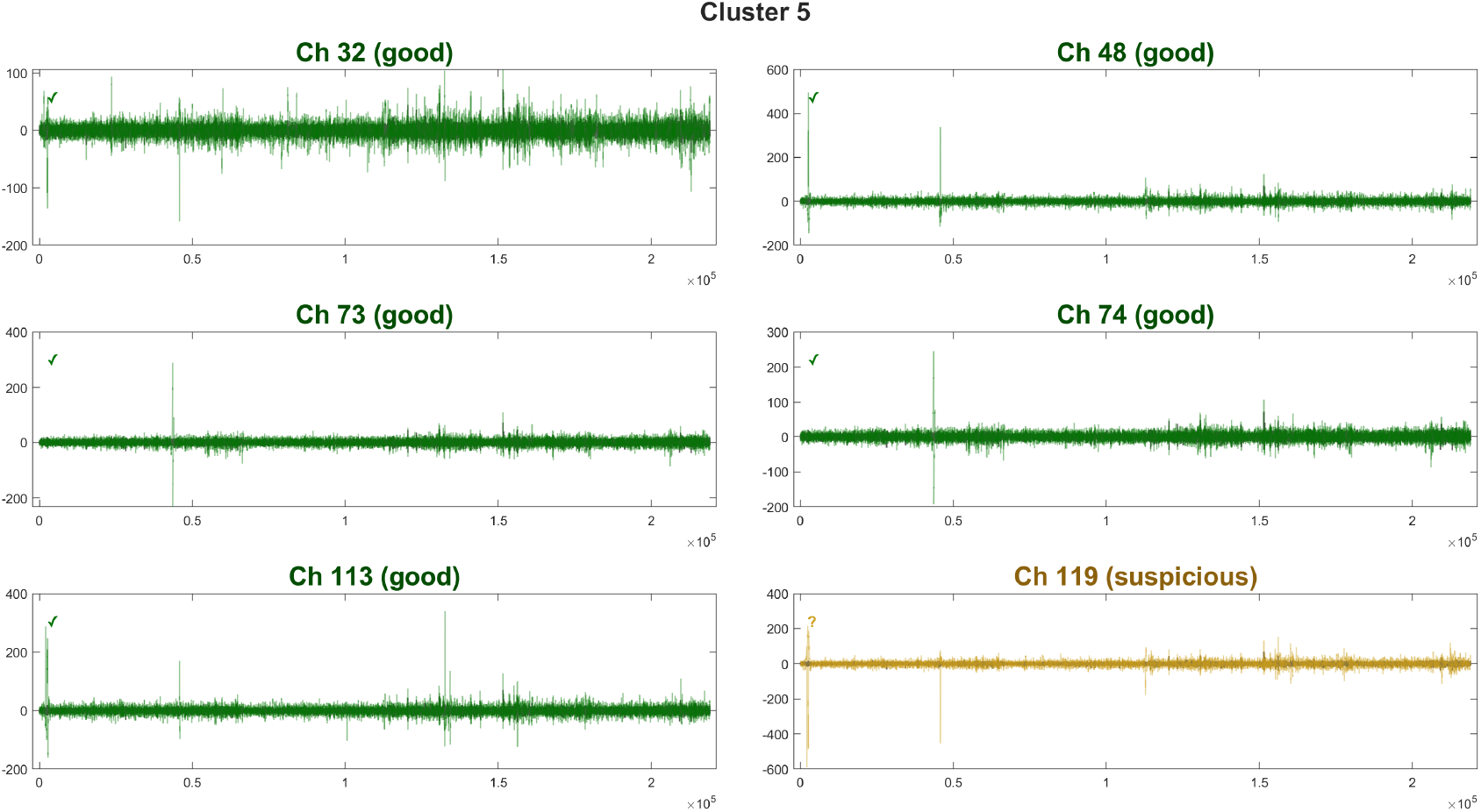
Page 6, cluster 5. We consider these channels to be acceptable. These artifacts are likely to be removed using ICA or other downstream means. None of these channels have strong flags, other than 119 being dissimilar with its neighbor in a small time frame.

**Fig. S8.**
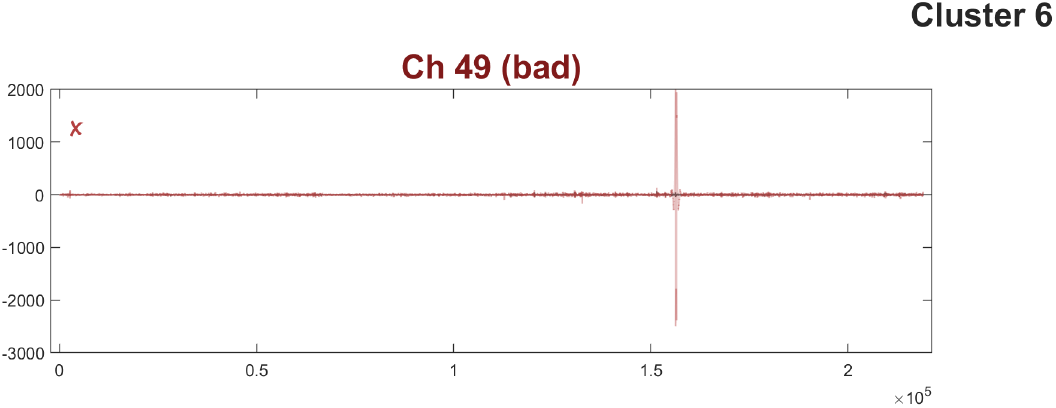
Page 7, cluster 6 (singleton). This channel was successfully tagged as *bad*. The transient around time sample 1.56 *×* 10^5^ is large enough that, in our experience, it would negatively impact subsequent ICA performance. One thing to keep it mind is that the neighbor dissimilarity is low; therefore, the artifact is likely to be present only in a small time frame.

**Fig. S9.**
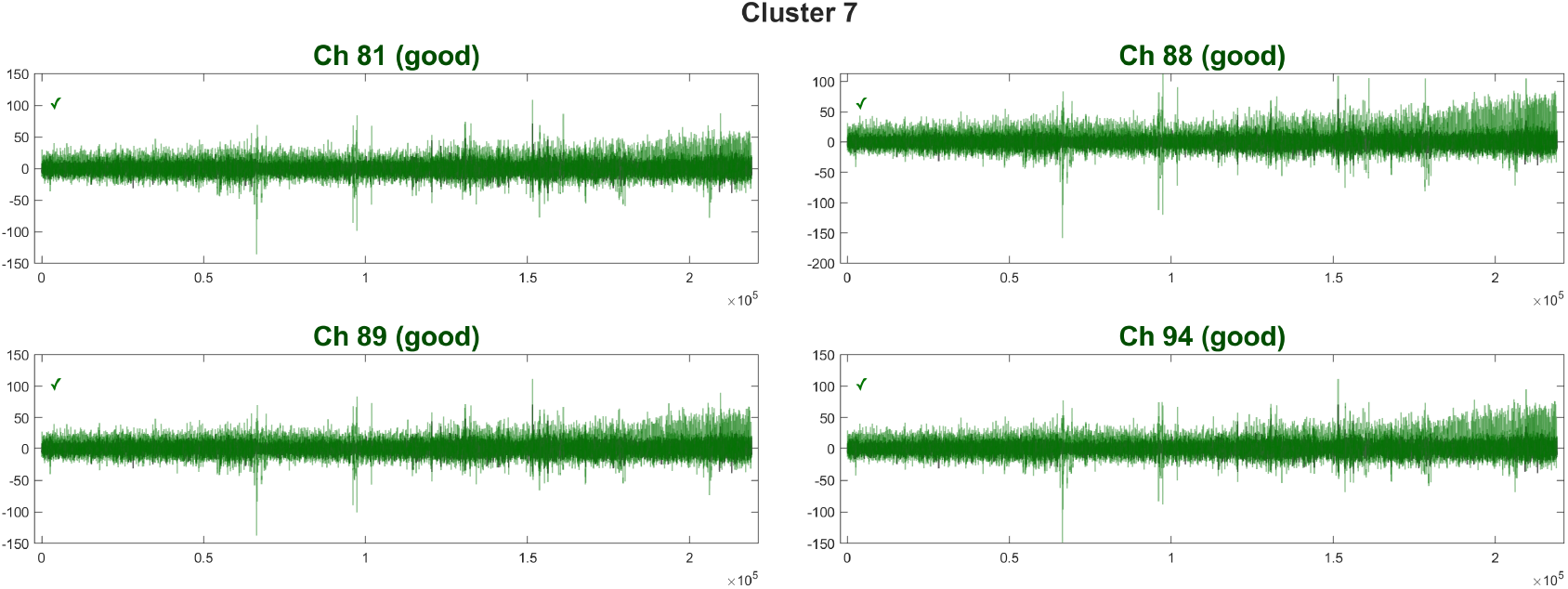
Page 8, cluster 7. We consider these channels to be acceptable. There are some unusual segments with increasing variance over time, which is likely the reason these channels were grouped together. These plots were not tagged as *suspicious* because they were not flagged by the general checks; however, the cluster size is small and therefore the UI shows this cluster.

**Fig. S10.**
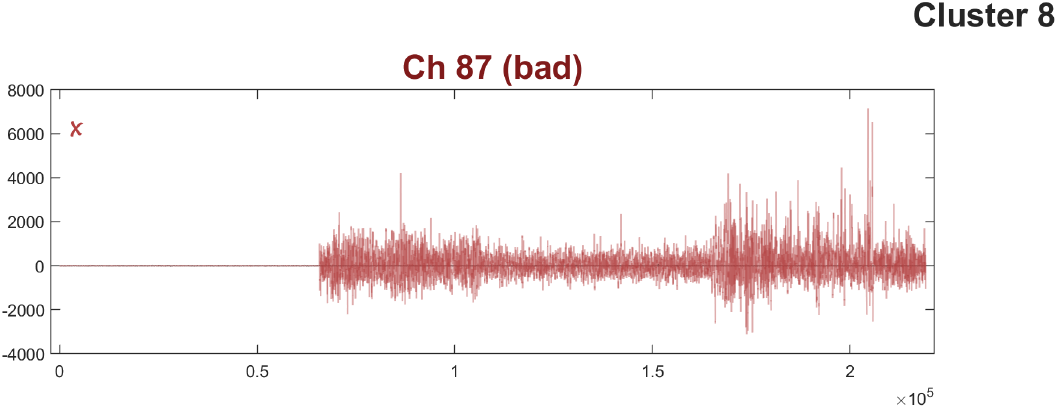
Page 9, cluster 8 (singleton). This channel was successfully tagged as *bad*. The amplitude and morphology of the data suggest that this channel may have lost connection to the scalp partway through the recording.

**Fig. S11.**
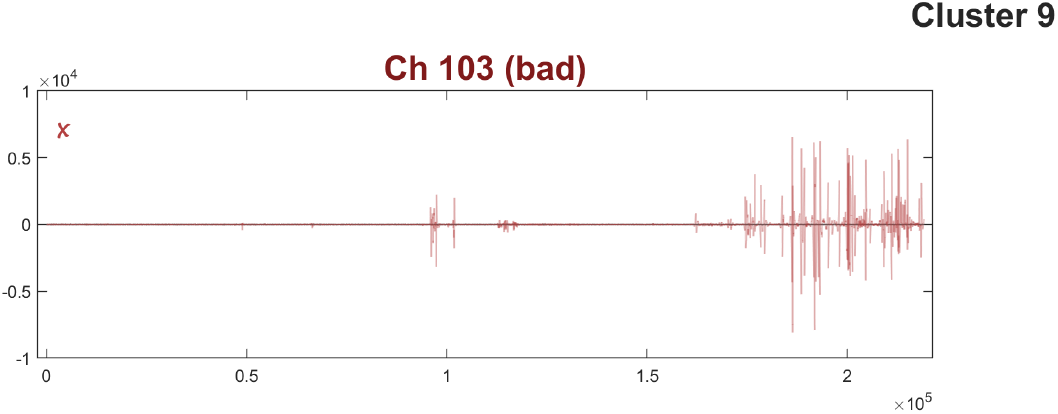
Page 10, cluster 9 (singleton). This channel was successfully tagged as *bad*. This channel is marked by intermittent, high-amplitude transients and should be removed.

**Fig. S12.**
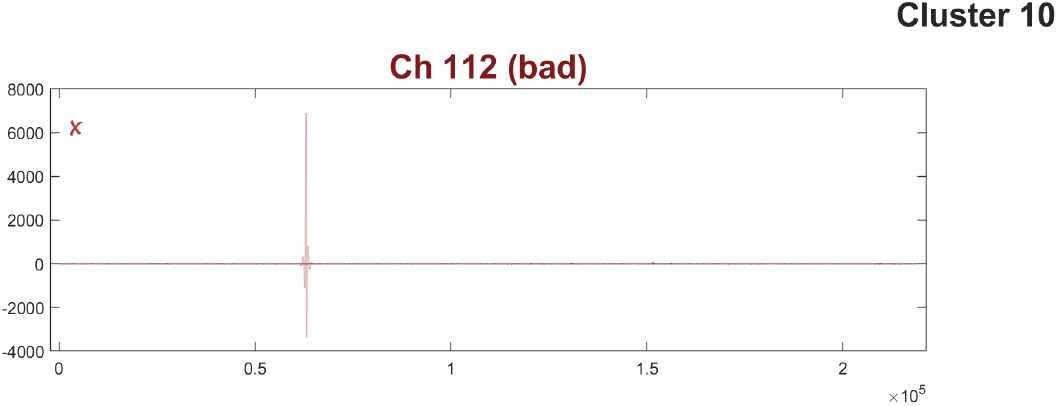
Page 11, cluster 10 (singleton). This channel was successfully tagged as *bad*. The transient around sample 1.56 *×* 10^5^ is too large and will affect ICA. Like Channel 49 (Figure S8), channel 112 has low neighbor dissimilarity and temporal variance equivalent to baseline. Therefore, the artifact is likely to be present only in a small time frame. In the future, this channel is a candidate for neighbor interpolation over the duration of the transient.

**Fig. S13.**
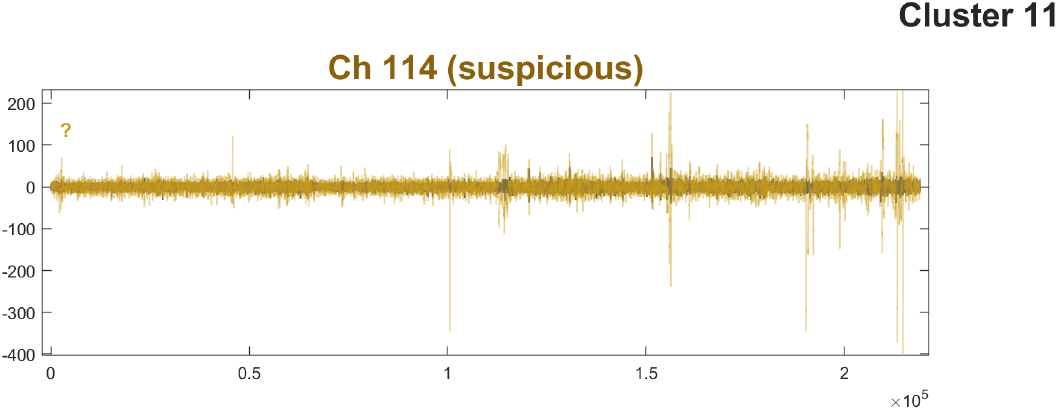
Page 12, cluster 11 (singleton). This channel is more challenging to characterize: this channel does not look too bad, but not good either. Two of the strong high-amplitude transients here are present in channel 91 (Figure S3) and channel 49 (Figure S8) — both of which were tagged for removal. Other transients found in channel 114 do not appear to repeat in other channels, but do not approach the amplitude-flag threshold. Looking at Figure 5, we can see that the suspiciousness scores were low, with the neighbor dissimilarity, max amplitude, and variance close to baseline values. Therefore, we would elect to leave this channel in. It will probably need some inspection after ICA and may require interpolation during the transient segments.

## Footnotes

1 https://github.com/edneuro/SENSI-EEG-Preproc-bad-ch

2 https://doi.org/10.25740/dg856vy8753

3 https://choosealicense.com/licenses/mit/

4 https://www.mathworks.com/products/matlab.html

5 https://www.mathworks.com/products/statistics.html

6 https://www.overleaf.com/latex/templates/henriqueslab-biorxiv-template/nyprsybwffws

